# Compensatory responses by managers, commercial and recreational harvesters to variation in stock abundance of Lake Erie walleye (*Sander vitreus vitreus*)

**DOI:** 10.1101/061143

**Authors:** Katrine Turgeon, Kevin B. Reid, John M. Fryxell, Thomas D. Nudds

**Author notes:** Co-authors.

## Abstract

Delayed quota adjustments, and/or lagged fishing effort and catch by harvesters, to changes in stock abundance may induce unstable population dynamics and exacerbate the risk of fishery collapse. We examined a 39-y time series of change to quotas by managers, and to effort and catch by both commercial harvesters and anglers, in response to changes in Lake Erie walleye abundance (*Sander vitreus*) estimated both contemporaneously and retrospectively. Quotas, commercial effort and catch were entrained by contemporaneous estimates of stock abundance. Recreational effort and harvest were not; they had better tracked abundance, as better estimated today, than did the commercial fishery. During the 1990s, a significant mismatch developed between the quota-driven commercial harvest and stock abundance that persisted until a new assessment process obtained. The quasi-open access recreational fishery, instead, freed anglers to respond better to stock abundance. Further elaboration of adaptive risk governance processes, including multi-model inference for stock assessments, may bode well to further reduce risk to fisheries imposed by lagged adjustments to variation in stock abundance.

## Introduction

Substantial theoretical effort has been devoted to the development of sophisticated models where resource abundance and harvest rate are dynamically coupled, and where uncertainty induced by environmental stochasticity, age structure, delayed life-history effects and density dependence is addressed (Lande et al. 1995, Punt and Hilborn 1997, Engen et al. 1997, Milner-Gulland et al. 2001, Jonzén et al. 2002, Beckerman et al. 2002, Coulson et al. 2008). Less attention has been devoted to managers’ and harvesters’ responses to changes in resource abundance in the short term, and the potential consequences for fisheries sustainability in the longer term (Botsford et al. 1983, Fryxell et al. 2010, Fulton et al. 2011). Accumulating empirical evidence and models suggest that weak or delayed compensatory responses by managers and harvesters to changes in resource abundance can induce cyclic or unstable population dynamics that could result in overfishing and increase the probability of a fishery becoming overfished (Bell et al. 1977, Botsford et al. 1983, Allen and McGlade 1986, Berryman 1991, Walters and Pearse 1996, Packer et al. 2009, Fryxell et al. 2010).

The outcome should depend on the degree of responsiveness by managers (through stock assessment and setting of annual quotas) and harvesters (through changes in fishing effort and harvest) to variation in stock abundance. For a fishery in which responses to variation in stock abundance are perfectly compensatory, rates of change in annual quota might be expected to be scaled to the magnitude and direction of expected rates of changes in stock abundance (Clark 1990, Walters and Pearse 1996). However, estimates of stock abundance used in stock assessments for purposes of establishing quotas may lack the precision or accuracy needed to detect changes in recruitment or natural mortality before stocks become overfished (Beamish and McFarlane 1983, Peterman and Bradford 1987, Walters and Maguire 1996). Managers attempting to balance ecological, economic and social objectives can face considerable uncertainty associated with stock assessments and model interpretation (Smith et al. 1999, Fulton et al. 2011). Similarly, responses by commercial and recreational harvesters to changes in stock abundance may also be expected to vary in magnitude and direction and to differ between the two types of harvesters. For example, management agencies often do not have the same control over effort and harvest of recreational fisheries as they do over quota-managed commercial fisheries (Post et al. 2012, MacKenzie and Cox 2013). Further, whereas recreational effort and harvest should be strongly driven by angler satisfaction (*i.e.*, angler catch rates), commercial harvesters should, so long as it is profitable, fish until quota is reached for a given year.

To examine responses by managers and harvesters to variation in stock abundance, we developed a conceptual framework with which to compare the magnitude and direction of annual rates of change in quota by managers, as well as effort and harvest by recreational and commercial harvesters, to annual rates of change in stock abundance. In this context, deviations from perfectly scaled fishery responses to variation in rates of change in stock abundance will vary with respect to the consequences they pose to the stability of the entire fishery (Fig. 1). A fishery more highly responsive to variation in stock abundance should exhibit (1) slopes of rates of change in quota, effort and harvest on rates of change in stock abundance close to 1; and (2) greater R^2,^ such that there are fewer, smaller deviations in directions that would, on one hand, increase the probability of overexploitation (positive deviations) and, on the other, increase the probability of poor economic or social outcomes for recreational (*i.e.*, low catch rates) and commercial fisheries (low catch rates, low quotas; negative deviation; Fig. 1).

**Figure 1.**
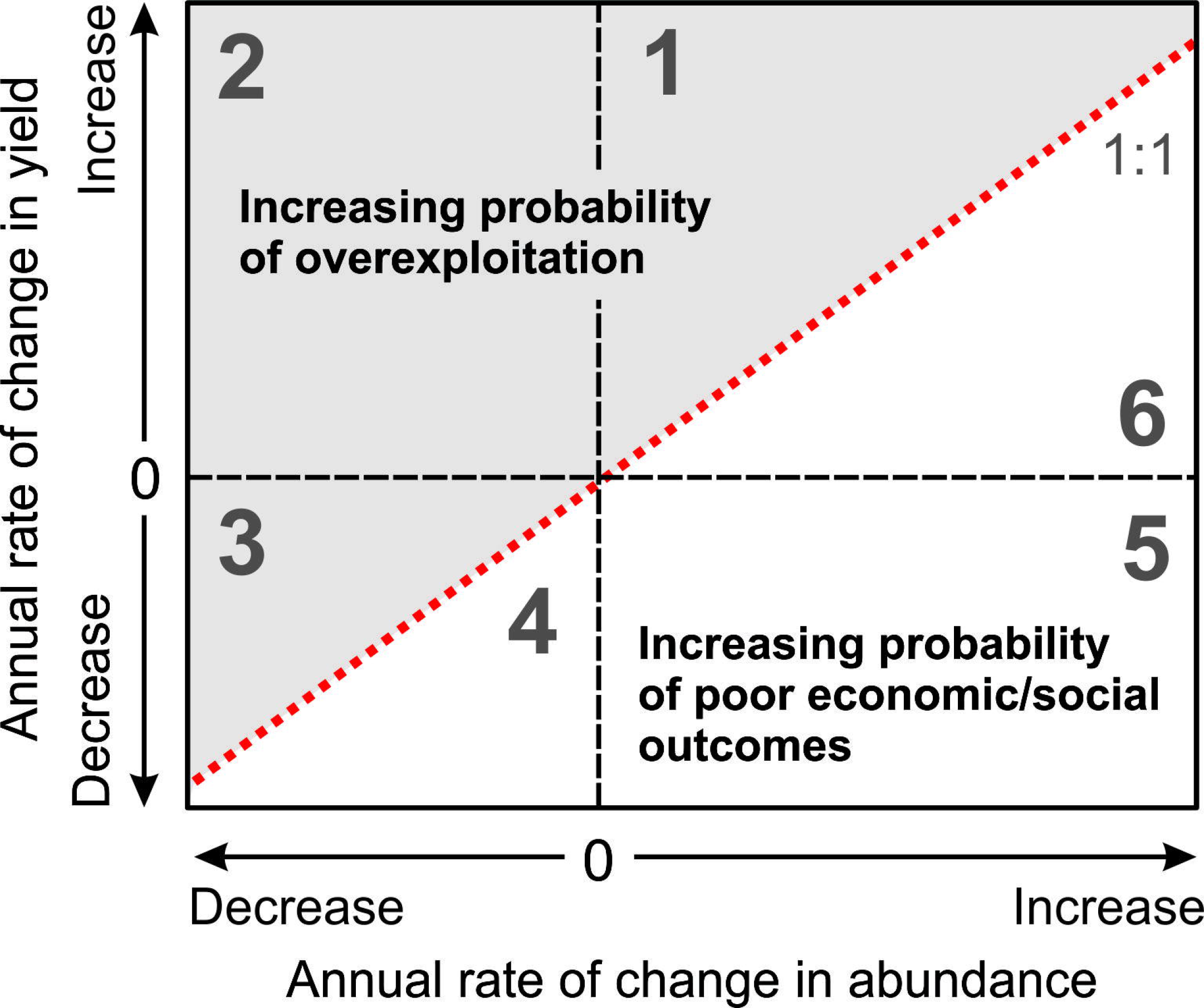
Conceptual framework for assessing compensatory adjustments by fisheries to annual rates of change in stock abundance by the magnitude and direction of deviations in annual rates of change in quota (and by extension, fishing effort or harvest). A fishery perfectly responding to variation in abundance should exhibit a slope of 1 (dotted red line). Above the dotted white line (positive deviations), changes in quota increase the probability of overexploitation; below it (negative deviations), changes to quota increase probability of poor economic or social outcomes for recreational and commercial fisheries (*i.e.*, opportunity costs, satisfaction). In regions 1 and 4, annual rates of change in quota overcompensate for changes in abundance, *i.e.*, they increase or decrease, respectively, faster than the rate at which abundance size changes. Similarly, in regions 3 and 6, quota responses undercompensate for changes in abundance. In regions 2 and 5, quota responses to changes in stock size diverged as to be opposing; in these regions, ecological risk to stocks and economic risk to fisheries, respectively, should be particularly exacerbated, corresponding to greater risk of fishery decline.

We used our framework and rich datasets for commercial and recreational walleye (*Sander vitreus vitreus*) fisheries in Lake Erie. Specifically, we examined to what degree annual rates of change in quotas set by managers, and corresponding annual rates of change in effort and harvest by commercial and recreational harvesters, tracked annual rates of change in estimates of walleye abundance available to managers at the time (*i.e.*, contemporaneous estimates) between 1975 and 2014. We also compared fishery responses to stock abundance retrospectively estimated from the statistical catch-at-age model (*i.e.*, retrospective estimates; stock catch at age-ADMB 2014) currently used by Lake Erie fisheries managers (Locke et al. 2005).

## Methods

### Study system

The Lake Erie walleye fishery is uniquely characterized by commercial and recreational fisheries that are largely confined to Canadian and to US waters, respectively, but share the same stock and management authority (Fig. 2), providing the opportunity to contrast between commercial and recreational fisheries controlling, to the extent possible, for potentially confounding sources of variation.

**Figure 2.**
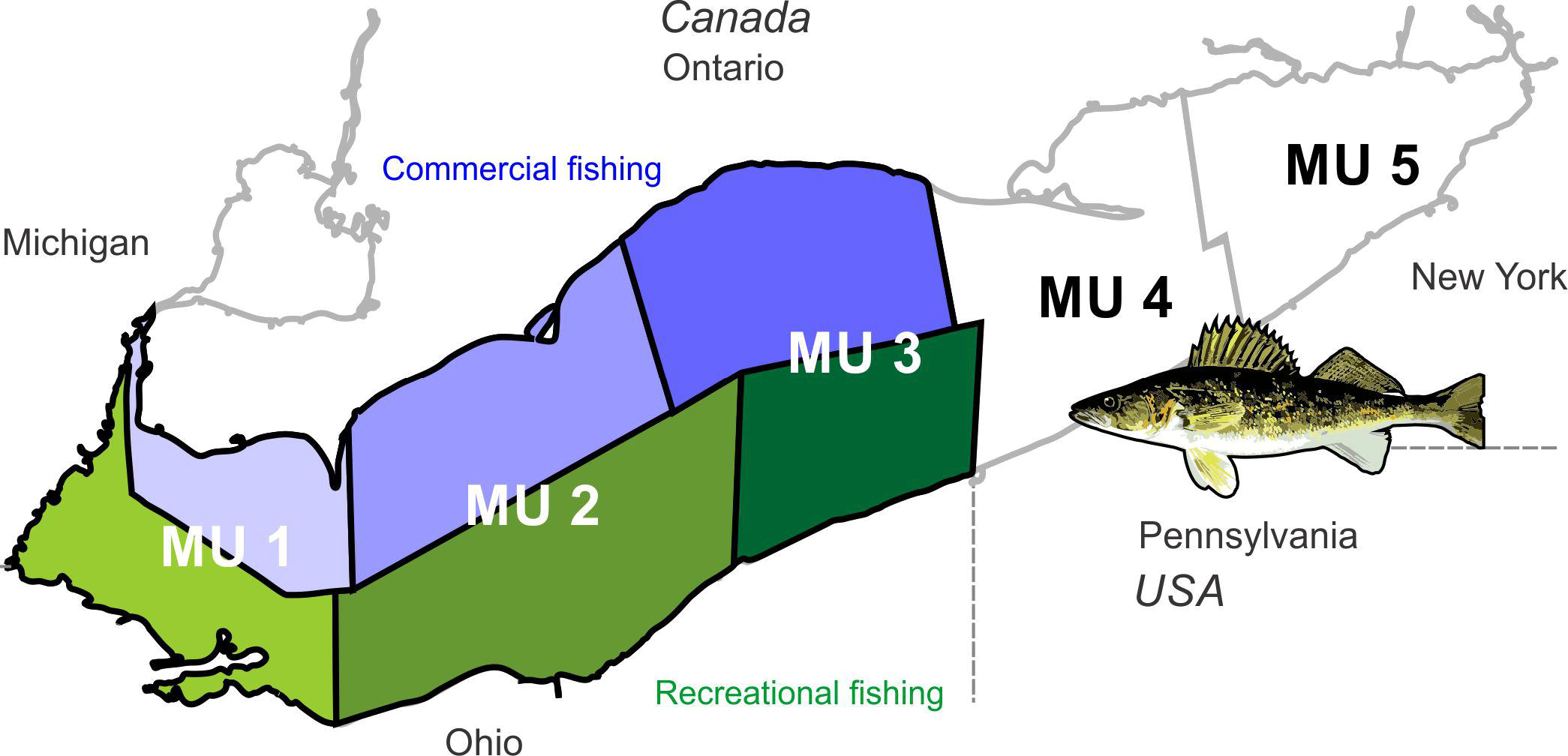
Five walleye management units (MUs) spanning west-central basins of Lake Erie.

In recent years, an average of about three million walleye were harvested annually by both commercial and recreational fisheries in Lake Erie (Lake Erie WTG 2015). The quota-regulated commercial fishery consists almost exclusively of a large mesh gill net fishery. The recreational fishery is officially assigned quota for purposes of allocating the annual total allowable catch (TAC) between it and the commercial fishery; in practice, it functions in a quasi-open access fashion. It consists of a charter boat industry, as well as boat- and shore-based individual anglers (and to a comparatively limited extent in Ontario; Brenden et al. 2012).

Lake Erie is the shallowest of the Laurentian Great Lakes, at a maximum depth of 64m, with surface area 25320 km^2^ and volume 473 km^3^. It comprises three natural basins and five walleye management units (MUs; Fig. 2). The western basin is most shallow (mean depth = 7.4 m; MU 1), biologically productive and the principal walleye spawning and nursery area (Nepszy 1977). The eastern basin (24.4 m; MUs 4 and 5) is least productive and the central basin (18.5 m; MUs 2 and 3) is intermediate (Ryan et al. 2003). The western walleye stock, the focus of the present analysis, spans mainly the west and central basins (MUs 1-3), the eastern stock spans MUs 4 and 5. Since 1978, western stocks consistently produced more than 95% of the total annual walleye harvest in Lake Erie (Lake Erie WTG 2015).

### Management of the Lake Erie walleye fishery

The Lake Erie Committee (LEC) comprised of members from each of four U.S. states and Ontario, Canada, proportionally allocates the TAC as quotas to the commercial and recreational fisheries based on recommendations from the Lake Erie Standing Technical Committee and the Walleye Task Group (Locke et al. 2005). Walleye stock abundance is cooperatively assessed on an annual basis using fishery-dependent and fishery-independent data and a statistical catch at age analysis (SCAA; (Lake Erie WTG 2015). A 39-y (1975-2014) time series of estimated walleye stock abundance, TAC (hereinafter quota), and fishing effort and harvest were available for both the commercial and recreational walleye fisheries from annual reports of the Walleye Task Group (http://www.glfc.org/lakecom/lec/WTG.htm#pub). Detailed histories of methods to estimate walleye stock abundance, determine quotas, and estimate commercial and recreational fishing effort and harvest are included in Appendix A.

### Data Analysis

To compare responses by managers (*i.e.*, annual rate of changes in quota), commercial and recreational fisheries (*i.e.*, changes in effort and harvest) to changes in estimated walleye abundance, we used generalized linear models (GLS) to regress annual rates of change in quota, commercial and recreational effort and harvest against the annual rate of change in abundance (*e.g.*, ln N_t+1_/N_t_). For these relationships, we tested whether slopes of regressions forced through the origin differed from 1 (perfect compensatory response to change in stock abundance, see framework in Fig. 1) using the following format: (y-x) ~ x where x = rate of change in abundance and y = rate of change in quota or harvest. We extracted the regression slope estimate and standard error of the relationships and calculated a pseudo-R^2^ using the deviance explained to get an estimate of the dispersion of the residuals. We used an autoregressive correlation structure (corAR1) to control for temporal autocorrelation in the GLS.We determined the autoregressive process in each time series by plotting each time series and by observing the autocorrelation function (ACF) and the partial autocorrelation function (PACF) on detrended data using an ARIMA (autoregressive integrated moving average model) diagnostic (astsa package v. 1.3 in R; Stoffer 2014). For each time series, errors were specified to follow an autoregressive process of degree 1 (*i.e.* the autocorrelation is highest between sequential years). GLS models with autocorrelated structure always had stronger AICc support (Burnham and Anderson 2002) than models without autocorrelated structure. We present only GLS models with the autocorrelated structure.

## Results

### General trends in walleye abundance, quota, effort and harvest

Between 1975 and 2014, estimates of walleye abundance, quotas, effort and harvest for commercial and recreational fisheries fluctuated widely (Table 1, Fig. 3). Quota adjustments by managers followed changes in stock abundance estimated at the time (contemporaneous estimates) quite well, both in magnitude and direction (Fig. 3 a). Commercial effort and harvest, tracked changes in quota and thus contemporaneous estimates in abundance (Fig. 3 a-b), whereas recreational effort and harvest did not (Fig. 3 a-c). Instead, recreational effort and harvest corresponded more closely with changes in abundance as derived from the latest catch-at-age model (Fig. 3).

**Figure 3.**
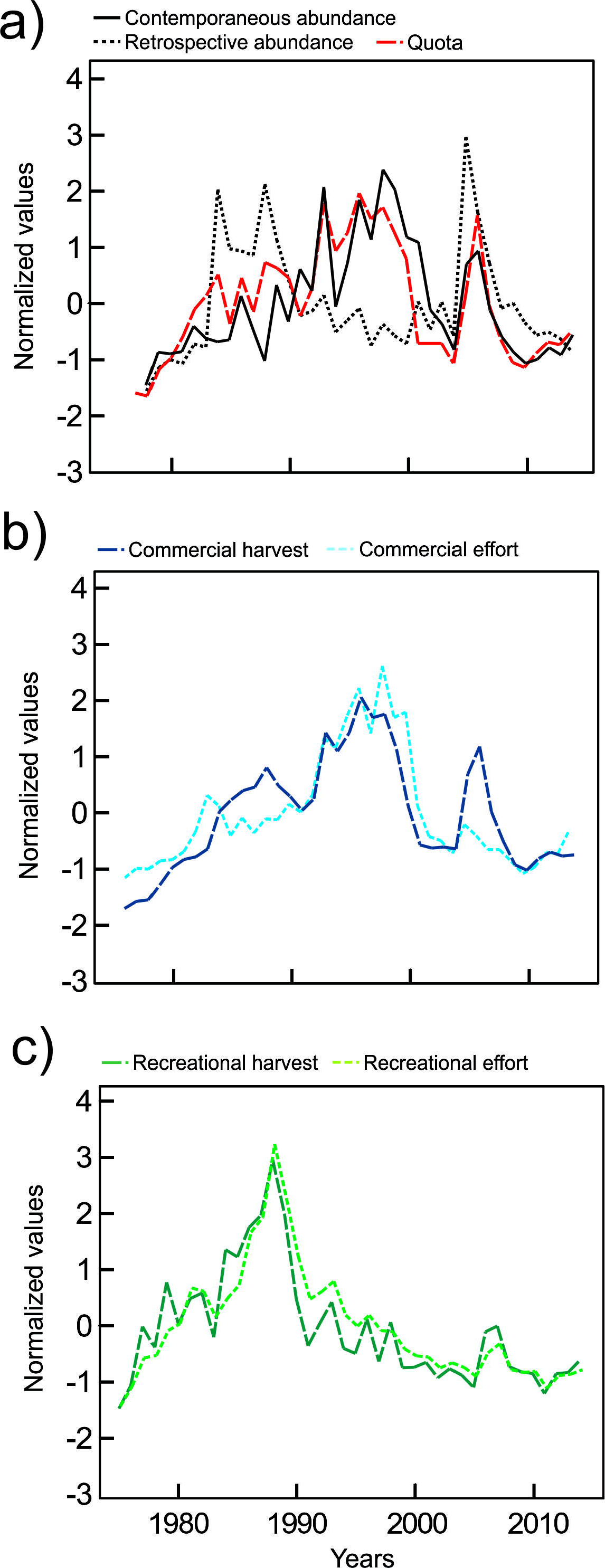
Time series (1975-2014) of a) walleye abundance contemporaneously estimated at year t (solid line; projected abundance extracted from the SCAA model used at that time), abundance retrospectively estimated in 2014 (solid line; using the latest SCAA-ADMB model 2014) and quotas (red dashed line), b) commercial effort (kms of gill net/year) and harvest and c) and recreational effort (angler hours/year) and harvest. Data were normalized. See Appendix A for details about SCAA models used.

**Table 1.** Mean, standard deviation (SD) and range of variables used to describe the dynamics of commercial and recreational walleye fisheries on Lake Erie between 1975 and 2014. Estimated walleye abundances were extracted from simulations of the SCAA used at the time of determining quota at year t (contemporaneous estimates) and from the most recent SCAA-ADMB model (2014; retrospective estimates). Values for abundance and quotas are in millions of walleye. See Appendix A for details about the SCAA models used for projecting contemporaneous and retrospective abundances.

| **Variable** | **Mean ± SD** | **Range (Min – Max)** |
| --- | --- | --- |
| Contemporaneous walleye abundance | $31.7 \pm 15.5$ | 9.46 – 68.2 |
| Retrospective walleye abundance | $45.5 \pm 23.7$ | 8.57 – 115 |
| Commercial Quota | $2.36 \pm 1.23$ | 0.32 – 4.76 |
| Recreational Quota | $3.24 \pm 1.60$ | 0.51 – 6.23 |
| Commercial harvest | $2.06 \pm 1.15$ | 0.16 – 4.47 |
| Recreational harvest | $2.26 \pm 1.54$ | 0.98 – 6.91 |
| Commercial effort (1000 kms of gill net/year) | $18.4 \pm 12.7$ | 3.73 – 51.6 |
| Recreational effort (1000 angler hours/year) | $4.96 \pm 3.04$ | 0.62 – 14.7 |

### Direction and magnitude of compensatory responses

Overall, annual rates of change in quotas, effort and harvest for both fisheries were positively correlated with annual rates of change in abundance (Fig. 4), but the slope of all regressed variables differed significantly from hypothetical values 1. The strength of the relation between rate of change in effort and harvest and the rate of change in abundance differed between commercial and recreational fisheries (Fig. 4). The annual rate of the change in quotas matched the rate of change in abundance (slopes = 0.71 and 0.73; R^2^= 0.28 and 0.32 for commercial and recreational fisheries, respectively; Fig. 4) Recreational harvest was more strongly (*i.e.*, higher slope), though more variably (*i.e.*, lower R^2^), compensatory to changes in abundance than was commercial harvest (recreational: slope = 0.31; R^2^ = 0.03, commercial: slope = 0.27; R^2^ = 0.25; Fig. 4). Changes in commercial effort and harvest were more strongly entrained by change in quotas (slopes, 0.51-0.57; R^2^= 0.50-0.69) than were changes in recreational effort and harvest (slopes, 0.22-0.39; R^2^, 0.10-0.11; Fig. 4). Harvest was strongly entrained by effort in both fisheries (Fig. 4).

**Figure 4.**
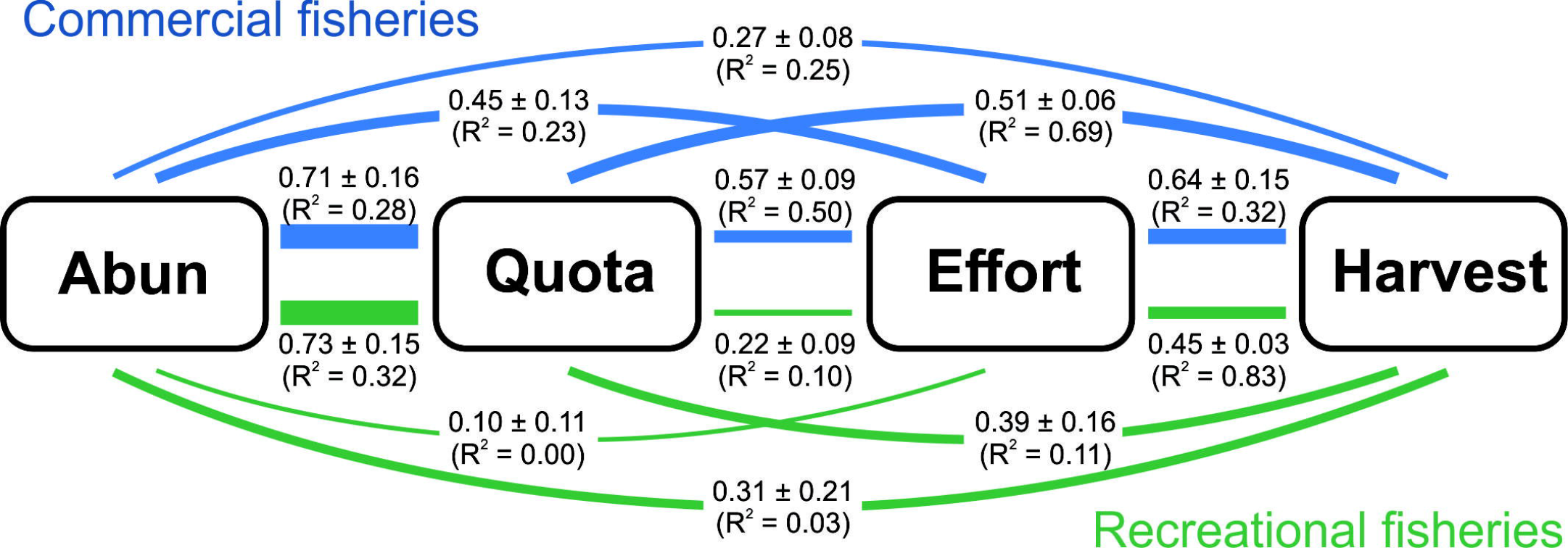
Estimate and standard error of the regression slope between contemporaneous abundance, quota, effort and harvest and pseudo R^2^ calculated to estimate the goodness-of-fit of the relationships for commercial and recreational Lake Erie walleye fisheries. Estimates are extracted from GLS with an autoregressive correlation structure (corAR1).

Deviations between the annual rate of change in quota and contemporaneous abundance estimates were relatively evenly distributed around the 1:1 relationship, *i.e.*, between increased probability of overexploitation (positive deviations) and increased probability of low or negative profits for commercial harvesters and poor angler satisfaction in recreational fisheries (negative deviations; Fig. 5 a). The magnitudes of the deviations from the 1:1 line were greater during the late 1980s and early 1990s and of much smaller magnitude since 2007, suggesting greater responsiveness, of improved magnitude and direction, to change in stock abundance (Fig. 6 a). Commercial harvest was strongly entrained by quota, so the direction and magnitude of the deviations between commercial harvest and abundance are comparable to those of quota (Fig. 5 b, Fig. 6 b). In recreational harvest, 56% of the deviations were in the regions corresponding with increased probability of overfishing (Fig. 5 c) and, contrary to quota and commercial harvest, the deviations in recent years were of high magnitude suggesting that the recreational fishery may be becoming less responsive to changes in abundance (Fig. 6 c).

**Figure 5.**
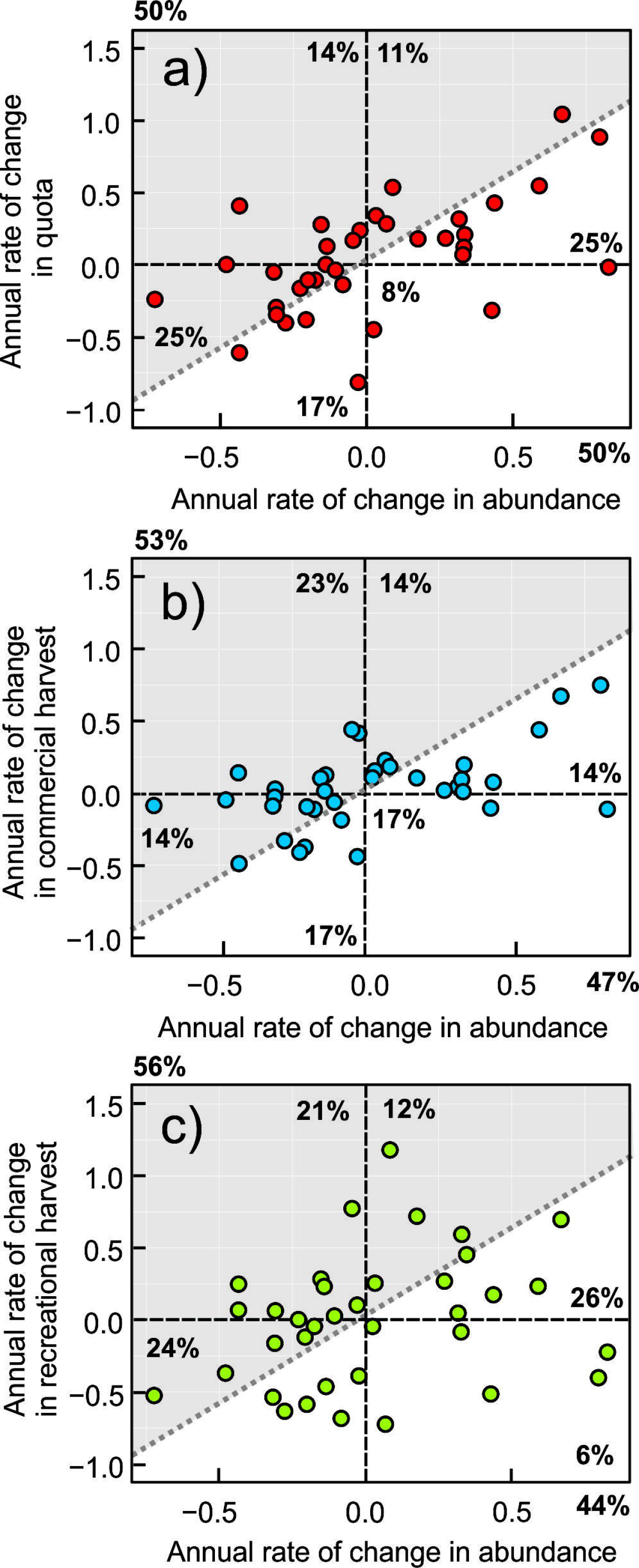
Observed annual rate of change in Lake Erie walleye contemporaneous abundance in relation to a) annual rate of change in quota, b) annual rate of change in commercial harvest and c) annual rate of change in recreational harvest. The grey dotted line represents a fishery perfectly synchronized with stock dynamics (see Fig. 1; conceptual framework).

**Figure 6.**
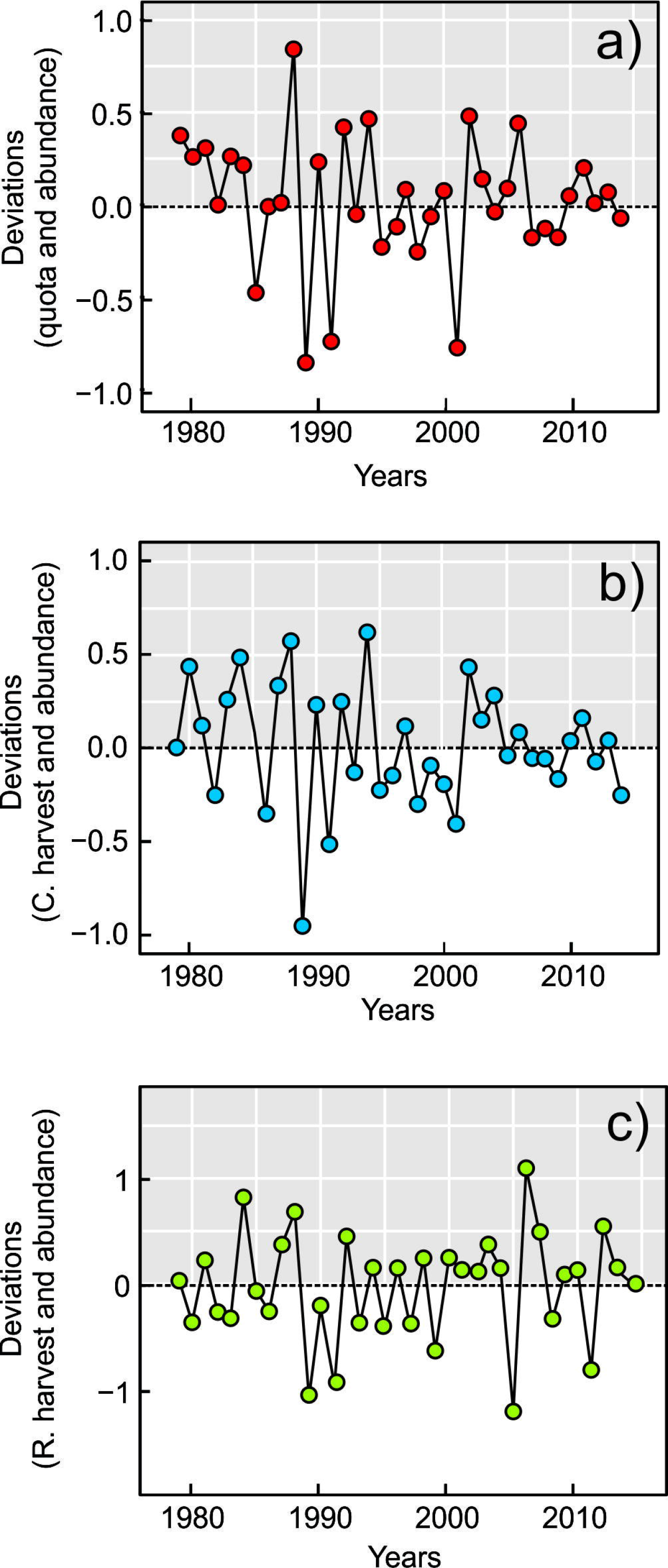
Deviations from perfect correspondence between the annual rates of change in contemporaneous abundance and the annual rate of change in a) quota, b) commercial harvest and c) recreational harvest between 1975 and 2014.

## Discussion

We developed a conceptual framework to evaluate compensatory responses of managers and harvesters to changes in stock abundance and used it to examine a rich dataset from Lake Erie walleye commercial and recreational fisheries. Managers were responsive in changing and setting quotas in response to available information about changes in stock abundance at the time (contemporaneous estimates). Commercial fisheries responded differently to changes in stock abundance and quotas than recreational fisheries. Commercial harvest and effort were strongly entrained by quotas whereas recreational fisheries better tracked abundance as retrospectively estimated.

There was an important discrepancy between contemporaneous abundance estimates and retrospective abundance estimates, arising in the early 1990s and until about 2000. Because commercial harvest and effort were strongly regulated by quotas in response to available information about stock abundance at the time, a mismatch developed between quota-driven commercial harvest and actual stock abundance. Specifically, quotas increased while stocks were declining, potentially generating the decline, observed today, in walleye abundance between the early 1990s and 2000.

### Disentangling decision making process from harvesting decisions

Accurate stock abundance estimates are essential to the setting of meaningful population abundance reference points and fishing quotas (Hilborn and Walters 2001, Hilborn 2002). Quota decisions were well reasoned at the time, though they were considerably vested in a single stock abundance model that, in retrospect, was unreliable (Locke et al. 2005) and led to a series of deviations in rates of change of quota in relation to rates of change in actual stock abundance that increased ecological risk (Fig. B1 a-b). Quotas entrained commercial harvest, resulting in harvest rates in excess of F=0.40 (Locke et al., 2005) and consequently a high probability of overfishing (Jiao et al. 2010). In the compensatory framework, the number of deviations between the annual rate of change in commercial harvest and abundance in the regions corresponding with increasing probability of overfishing increased from 53% to 65% (Fig. B1 c-d). Thus, a series of small, consecutive, unscaled and untimely compensatory adjustments of quotas in response to changes in walleye abundance generated potentially long-term social, economic and ecological negative impacts. These unscaled adjustments were apparently due to a false impression of a healthy walleye stock at that time.

After 2003, Lake Erie walleye fisheries managers adopted a state-dependent harvest control rule which allowed a more nimble management response to changes in resource abundance (see Locke et al. (2005) for details). This management rule obliged managers to rapidly increase quotas in 2005 and 2006 following the recruitment of the strong 2003 year class, and to gradually decrease them during the 2007-2011 period as that exceptionally strong year class progressed through the fishery (corresponding to smaller magnitudes of deviations between annual rate of change in abundance and quota). Nonetheless, deviation using retrospective estimates in abundance suggest that quota responses continue to periodically strongly over-and under-compensate for changes in estimated stock abundance (Fig. B1 b), consistent with the idea that model uncertainty still manifests and might lend to lagged or weak responses to changing stock abundance, exacerbating the likelihood of negative outcomes for the entire fishery.

By trusting erroneous projections from the CAGEAN model, managers were not aware of a decrease in population, even though some indicators of stock status (*i.e.*, increase in commercial effort, decrease in commercial CPUE, general decrease in effort and recreational harvest) indicated different trends from model estimates. This situation is similar to what happened in the Icelandic cod fishery in late 1990s where, following an appearance of a healthy cod stock (*i.e.*, increases in catches and high estimates from stock assessment models), managers raised the quotas to capitalize on the apparent increase. When they discovered that stock was overestimated, they drastically reduced quotas, causing severe economic losses to the fishery (Rosenberg 2003). Similar erroneous projections from incorrect estimates of natural mortality or fish age structure in the Pacific Ocean perch (*Sebastes alutus*; Canada) in the orange roughy (*Hoplostethus atlanticus*, New Zealand) and in the Northern cod (*Gadus morhua*; Canada) are suspected to be contributing factors in the failure of these fisheries (Beamish and McFarlane 1983, Mace et al. 1990, Walters and Maguire 1996, Myers et al. 1997).

### Reasons for different responses of the two fisheries

To the best of our knowledge, no study has compared responses of commercial and recreational fisheries to changes in stock abundance and quota. Other studies comparing recreational and commercial fisheries contrasted the characteristics, short-term impacts and behaviours of commercial and recreational fisheries in relation to conservation and management (*e.g.* by-catch, trophic changes, fisheries induced selection, management strategies; Murray-Jones & Steffe 2000; Policansky 2001; Cooke & Cowx 2004, 2006; Sutter *et al.* 2012; Cooke & Murchie 2013). Detailed information on the responses of recreational anglers to changing resource abundance is rare, particularly recreational catch in relation to effort (Post *et al.* 2002; Cooke & Cowx 2004).

The observed differences in the responsiveness of the two fisheries may be related to respective motivations of recreational and commercial harvesters, from social pressures and degrees of confidence in stock assessments from a manager’s perspective. In either case, outcomes depend on the degree of responsiveness of harvesters and managers to change in stock abundance. If harvesters’ effort changes rapidly in response to changing resource abundance, harvest should simply track variation in stock abundance induced by environmental variation. We attribute this rapid adjustment of effort by anglers to the large, highly-networked charter boat fleet and angling groups in Ohio and Michigan, to numerous online resources about the status of the walleye fishery for anglers, and intense coverage of the fishery in print media and other news outlets. Social sharing of information about the state and quality of these fisheries is likely very rapid in this particular valuable fishery system. However, recreational harvest and effort was less responsive to more recent rapid changes in quota and abundance (Fig. 3 b) as indicated by higher magnitude deviations after 2000 (lower R^2^; Fig. 6 c and Fig. B1 e-f). This reduced responsiveness may be attributable to ongoing changes in angler demographics. Recruitment of new anglers is not compensating for the older anglers that are leaving the freshwater recreational fishery, especially in the U.S. (The Outdoor Foundation 2014), but also in Canada where the total number of days spent angling had been in decline since the mid-1990 (Hofmann 2008, DFO 2012). In Lake Erie, the reduced fishing power of the recreational fishery, possibly in combination with reduced rates of communication and networking among fewer and younger anglers, appears to constrain the responsiveness of the recreational fishery.

On the other hand, our analysis suggests that the commercial walleye fishery was strongly regulated by annual variation in quotas. As long as it is profitable, commercial fishermen have a strong incentive to continue fishing for walleye until the quota is reached, but, unlike recreational anglers, they will tend to stay in the fishery because of significant investment in quota, vessels and gear, and because many commercial fishermen have options to gain alternative revenues by fishing for species other than walleye. Management of a declining fish stock is typically a delay-ridden process. Because catch reductions impose short-term losses on fishers, merchants, and processors, managers face political opposition—and even legal challenges—and they may hesitate to change the *status quo* (Rosenberg 2003).

### Management, harvesting decisions and stock stability

Several empirical and theoretical studies have suggested that delayed harvesting responses can contribute to fluctuations in stock abundance (Bell et al. 1977, Botsford et al. 1983, Allen and McGlade 1986, Berryman 1991, Fryxell et al. 2010, Nilsen et al. 2011). Harvest data on whitefish (*Coregonus clupeaformis*) in Lesser Slave Lake (Alberta) suggest cyclic fluctuations in whitefish abundance driven by a lagged relationship between fishing effort (the number of licenses issued) and fish abundance (catch per licence; Bell et al. 1977). Fryxell et al. (2010) suggested that delayed changes in hunting effort and quotas contributed to quasi-cyclic fluctuations in populations of white-tailed deer (*Odocoileus virginianus*) in Ontario and moose (*Alces alces*) in Norway. Two studies of the northern California dungeness crab (*Cancer magister*) fishery suggested that delayed entry or departure from the crab fishery and lags in market expansion and contraction following changes in abundance contributed to crab population cycles (Botsford et al. 1983, Berryman 1991). A simulation study (McGarvey 1994) applied to Georges Bank sea scallop (*Placopecten magellanicus*) fishery suggested that density-dependent recruitment and environmental variability combined with a dynamic harvest process dictated by variable effort could generate irregular cycles of stock abundance and effort. Simulations based on a dynamic model of a Nova Scotian groundfish fisheries suggest that lagged responses in fishing effort could amplify rapid fluctuations in haddock (*Melanogrammus aeglefinus*; Allen and McGlade 1986). Delays in the response of commercial fishing effort and quotas to changes in walleye stock abundance could have contributed similarly to fluctuations over time.

In contrast, effort by recreational walleye anglers changed quickly in response to changing walleye stock abundance, perhaps due to changes in fishing satisfaction (Smith 1999, Post et al. 2002), resulting in a better match between exploitation rates and fish stocks than in the commercial fishery (Johnson and Carpenter 1994, Hilborn and Walters 2001, Fryxell et al. 2010). In this case, the annual harvest data for the walleye commercial fishery seem less reliable as an indicator of stock status than are the same data from the recreational fishery, because the latter was seemingly more responsive to biological conditions. These results reinforce the potential utility of long term time series data from both commercial and recreational fisheries to understand fish population dynamics.

Finally, our results suggest that continued improvement in fishery responses to changes in stock abundance, to minimize both ecological and economic risk, will involve a better appreciation, and exploitation, of knowledge about the dynamical relationships among fish stocks, managers and fishermen. Success in fishery management lies not only in understanding ecological uncertainty arising from stock dynamics, but also the critical role and uncertainty coming from the human behaviour (Fulton et al. 2011). The success of fishery management relies not only on understanding and dealing with uncertainty arising from stock assessments but also on accounting for the critical role of human-induced uncertainties (Haapasaari et al. 2007, Fulton et al. 2011). Our results are consistent with the idea that explicit ecological modelling of manager and harvester behaviour in response to stock dynamics, and vice versa, should help to achieve more pragmatic management policies. In addition to seeking better understanding of the relationships among manager and harvester behaviour and fish population dynamics, a pragmatic governance system would eschew management reliance on any single stock assessment model; relying instead on an increasing role for interdisciplinarity, multimodel inference and other decision analytical tools to foster management decisions that are ecologically, economically and socially robust to both ecological and human-induced uncertainty.

## Acknowledgments

This work was supported by funding from the Natural Sciences and Engineering Research Council of Canada (NSERC) Discovery Grant Program to JMF (STPGP-2009-380926), NSERC Canadian Fisheries Research Network (CFRN; NETGP 389436-06) and Ontario Commercial Fisheries’ Association (OCFA) grants to TDN, and Fonds Québécois de la Nature et des Technologies (FQRNT) postdoctoral scholarship to KT. We thank R. Norris, F. Zhang, A. Debertin and B. Locke for helpful suggestions on an early draft of the manuscript.

